# singletCode: synthetic barcodes identify singlets in scRNA-seq datasets and evaluate doublet algorithms

**DOI:** 10.1101/2023.08.04.552078

**Authors:** Ziyang Zhang, Madeline E. Melzer, Karun Kiani, Yogesh Goyal

## Abstract

Single-cell RNA sequencing datasets comprise true single cells, or singlets, in addition to cells that coalesce during the protocol, or doublets. Identifying singlets with high fidelity in single-cell RNA sequencing is necessary to avoid false negative and false positive discoveries. Although several methodologies have been proposed to infer true singlets and doublets, they typically rely on datasets being highly heterogeneous. Here we develop and apply singletCode, a computational framework that leverages datasets with synthetically introduced DNA barcodes for a hitherto unexplored application: to extract ground truth singlets. We demonstrate the feasibility of singlets extracted via singletCode to evaluate the performance and robustness of existing doublet detection methods. We find that existing doublet detection methods are not as sensitive as expected when tested on doublets simulated from experimentally realistic ground truth singlets. As DNA barcoded datasets are being increasingly reported, singletCode can identify singlets and inform rational choice of doublet detecting algorithms and their associated limitations.

## Introduction

Rapid advances in single-cell RNA sequencing (scRNA-seq) technologies have enabled the characterization of cellular gene expression at an unprecedented resolution and scale. Such technologies have revealed extensive functional diversity of cell states across biological contexts, including cancer, evolution, and development. Briefly, scRNA-seq technologies rely on distributing individual cells from a suspension into individual reactions, each labeled with a unique “ID”, usually in the form of a reaction-specific sequence barcode. Despite numerous technological optimizations, multiple cells can occasionally be encapsulated in a single reaction, resulting in doublets or multiplets where two or more cells are assigned the same reaction ID (**Figure 1A**). The percentage of doublets in a given experiment depends on several factors, including the features of the sample and throughput, and can be as high as 40-50% [1,2]. In turn, such artifacts affect the downstream analysis [3]. Indeed, a central challenge in the burgeoning scRNA-seq field is to identify true single cells (“singlets” from here on) and therefore accurate samples of individual cells’ transcriptomes in the resultant scRNA-seq datasets.

**Figure 1.**
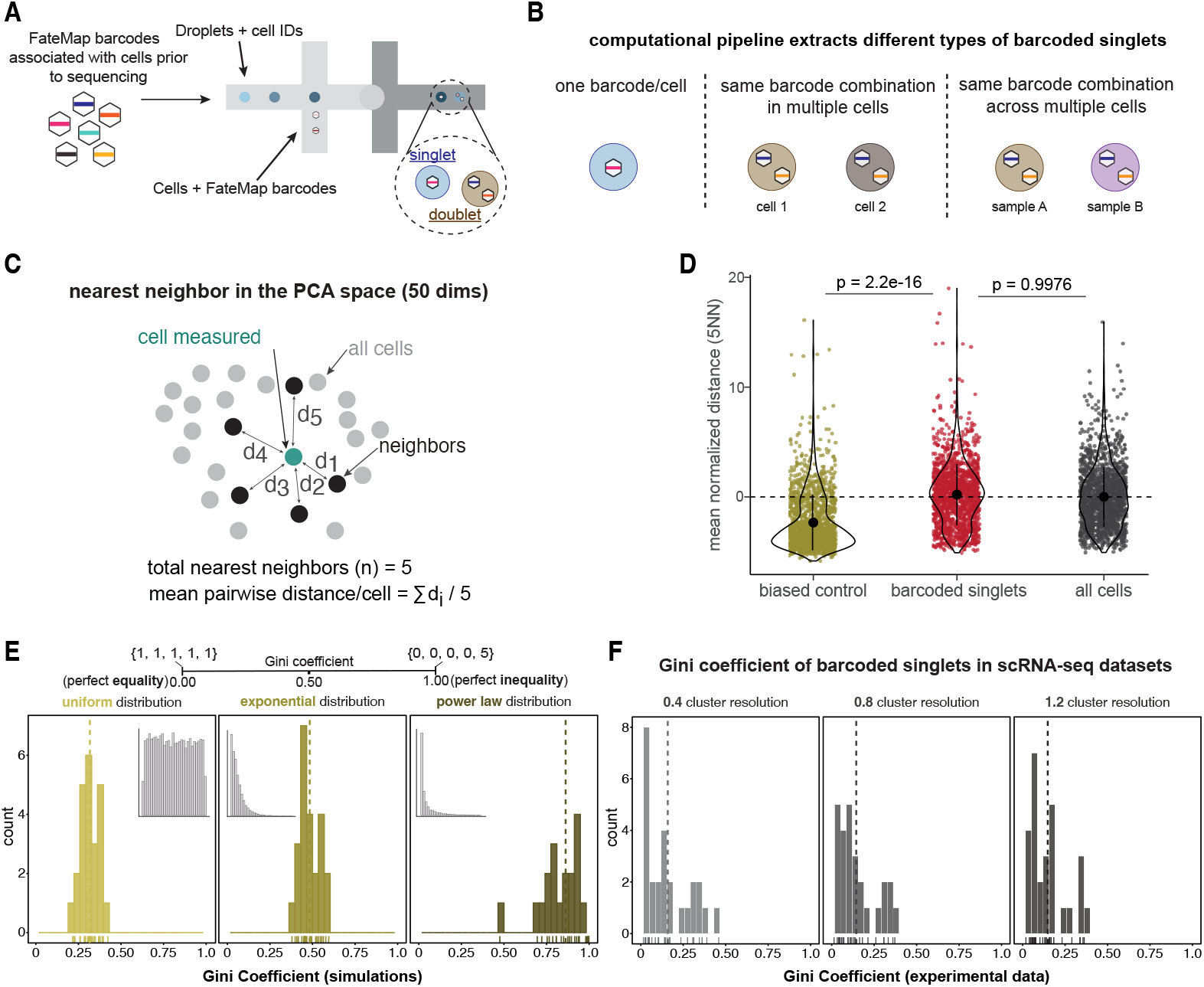
A. FateMap lineage barcodes exist in cells *prior* to the implementation of the single-cell sequencing protocol where droplets add a unique cell ID. A singlet is one droplet with a single cell such that there is a 1:1 mapping of a FateMap barcode to a cell ID. Alternatively, an example of a doublet is when one droplet has two or more cells with each having a unique barcode. B. The three definitions of singlets for the purposes of our analyses: (1) One droplet with one FateMap barcode and one cell; (2) the same FateMap barcode combination found in multiple cell IDs, suggesting the two cells are progeny of a single cell; (3) the same FateMap barcode found in cells across multiple “twin” samples suggesting the two cells are also progeny of a single cell. C. Schematic of the pairwise distances of a cell’s five nearest neighbors in the 50 dimensional principal component space. The mean distance for each cell is used to describe the degree of transcriptional similarity to other cells. D. Average nearest neighbor distances are plotted for three types of datasets: a biased negative control where cells are artificially clustered, the barcoded singlets, and all barcoded cells (singlets and multiplets). Each dot represents the average nearest neighbor distance of a cell. The p-values were obtained using a one-sample Wilcoxon rank sum test. E. (top) A schematic to demonstrate how different sets of numbers (braces) influence the values of Gini coefficient. A set of numbers with perfect equality results in a value of zero while a set of numbers with perfect inequality will lead to a value of one. (bottom) The distribution of simulated Gini coefficients (total ∼20 simulations) from randomly sampling three different distributions: uniform (mean 0.305), exponential (mean 0.529), and power (mean 0.829). Insets: histogram of values of the distribution being used for Gini calculations. Dotted lines: mean value. F. Histogram of distribution of Gini coefficient of proportion of singlets across all clusters for three different cluster resolutions. Resolution of 0.4 had a mean Gini coefficient across samples of 0.159, with a range of 0.015-0.459; Resolution of 0.8 had a mean Gini coefficient of 0.145 with a range of 0.014-0.371; Resolution of 1.2 had a mean Gini coefficient of 0.148 with a range of 0.019-0.363. Dotted line: mean value. See also Supplementary Figure 2.

Several computational frameworks have been developed to identify singlets in scRNA-seq datasets [1,2,4–10]. Although each framework deploys its own algorithm, such methods typically rely on gene expression differences between individual cells to remove cells with a putative mixture of different expression profiles. As such, the algorithms naturally necessitate datasets consisting of vastly different cell types or species, where doublets are collectively referred to as heterotypic doublets. Therefore, such methods do not work well with “transcriptionally similar” cells [2], where doublets are referred to as homotypic doublets. Between these two extremes is a scenario more representative of many experimental designs in which cells are not vastly different, and in cases where continuums of cell states within a particular cell type can have functional consequences. However, the performance of doublet detection methods has not been systematically investigated on such realistic datasets. Furthermore, such algorithms take inferred singlets as inputs and do not have “ground truth” singlets, posing further challenges in identifying *bona fide* singlets. Although certain experimental techniques such as cell hashing [11] or lipid tagging [9] can help increase the confidence in doublets identified, they do not necessarily provide unique identifiers at single-cell resolution.

Recent developments in DNA barcoding methodologies have added another dimension to scRNA-seq datasets, revealing unique transcriptional signatures and clonal dynamics [12–26]. Here, we describe our framework, singletCode, which leverages DNA barcoding for a new application: identifying “true” singlets in scRNA-seq datasets. We posited that since DNA barcoding allows for individual cells to have a unique identifier *prior* to scRNA-seq protocols, these barcodes could help identify “true” singlets. The singlet population can then be used to simulate “ground truth” doublets and compare performance of the doublet-finding algorithms. Our proof-of-concept analysis was implemented on 27 barcoded scRNA-seq datasets, covering 141,044 total cells across several cell types and experimental designs. We find that existing state-of-the-art doublet detection methods show lower than reported sensitivity to doublets simulated with ground truth singlets. Since the repertoire of such datasets is increasing rapidly and DNA barcoding is becoming commonplace, our framework provides rational guidance for identifying singlets and choosing appropriate doublet-finding algorithms.

## Results

Since various studies use different heuristics and thresholds for barcode assignment [26–29], we first developed a standardized pipeline that can reliably extract true singlets based on the synthetically introduced FateMap lineage barcodes (see **Methods**). FateMap barcodes are 100 base pairs long and offer up to ∼50-60 million library complexity [26], which ensures multiple packaging copies of the same barcode are rarely present in an experimental design by chance. In short, we label “singlets” as those cells that, as a result of multiplicity of infection, satisfy one of the three conditions: (1) a single barcode identified per cell ID; (2) multiple barcodes per cell but the same barcode combination is found in other cells in the same sample; and (3) multiple barcodes per cell but the same barcode combination is found in other cells across samples within the same experimental design (common for barcoding studies [15,26–28,30]) (**Figure 1B**). The recovery percentage of singlets with this standardized pipeline was 81.7% as compared to those previously reported (50-60%) for the same datasets [26]. Summarily, our pipeline identified barcoded singlets for 27 scRNA-seq datasets obtained from 9 experimental designs and a total of 141,044 cells [26–28]. Note that previous studies have established that the introduction of synthetic barcodes does not affect the ensemble of cell types in a population [26,28]. These datasets cover various cell types (two patient-derived melanoma cell lines, one triple negative breast cancer cell line, stem cells, a fibroblast-like cell line, an induced pluripotent stem cell line, and primary melanocytes), biological processes (drug resistance, differentiation, reprogramming), and technical sequencing specifications (**Supplementary Table 1**) [26–28]. We annotated singlets within each scRNA-seq dataset and simulated doublets by averaging the transcriptome of two true singlets (see **Methods**). Our curated datasets are thus the closest attempt to benchmark doublet detection methods with ground truth singlets and doublets.

We next asked whether barcoded singlets exhibited a preference for specific cell states, as a bias would limit the scope of our approach’s usability. First, we calculated the average distance of each singlet to its first five nearest neighbors in high dimensional principal component space (**Figure 1C**). We then compared the singlet average neighbor distances to those obtained from scRNA-seq dataset consisting of all cells (singlets and doublets) and those artificially subsampled to mimic bias (see **Methods**). We reasoned that if the average neighbor distances of singlets were indistinguishable from all cells, the singlets uniformly spanned the entire scRNA-seq dataset. Indeed, singlet nearest neighbor distances were indistinguishable from the entire scRNA-seq dataset and significantly different from the biased control dataset (**Figure 1D, Supplementary Figure 1**). Note that this analysis was independent of the cluster designation in the Uniform Manifold Approximation and Projection (UMAP) space. Second, we used the Gini coefficient, a metric used to measure inequality in populations (0 and 1 imply perfect equality and inequality, respectively; 0.33 for simulated uniform distributions) (**Figure 1E**) [31]. We calculated the Gini coefficient of the proportion of singlets in each UMAP cluster across all 27 scRNA-seq datasets. For a cluster resolution of 0.4, the mean Gini coefficient across datasets was 0.159 (range of 0.02-0.46), which was between perfect equality (mean = 0) and uniform distributions (mean = 0.305). Our results were robust across a wide range of shared nearest neighbor resolutions (0.4–1.2), which cause different total numbers of clusters on the UMAP (**Figure 1F, Supplementary Figure 2**). Collectively, our analyses do not suggest the presence of systematic bias toward particular cell types or clusters when identifying singlets through FateMap barcodes.

**Figure 2.**
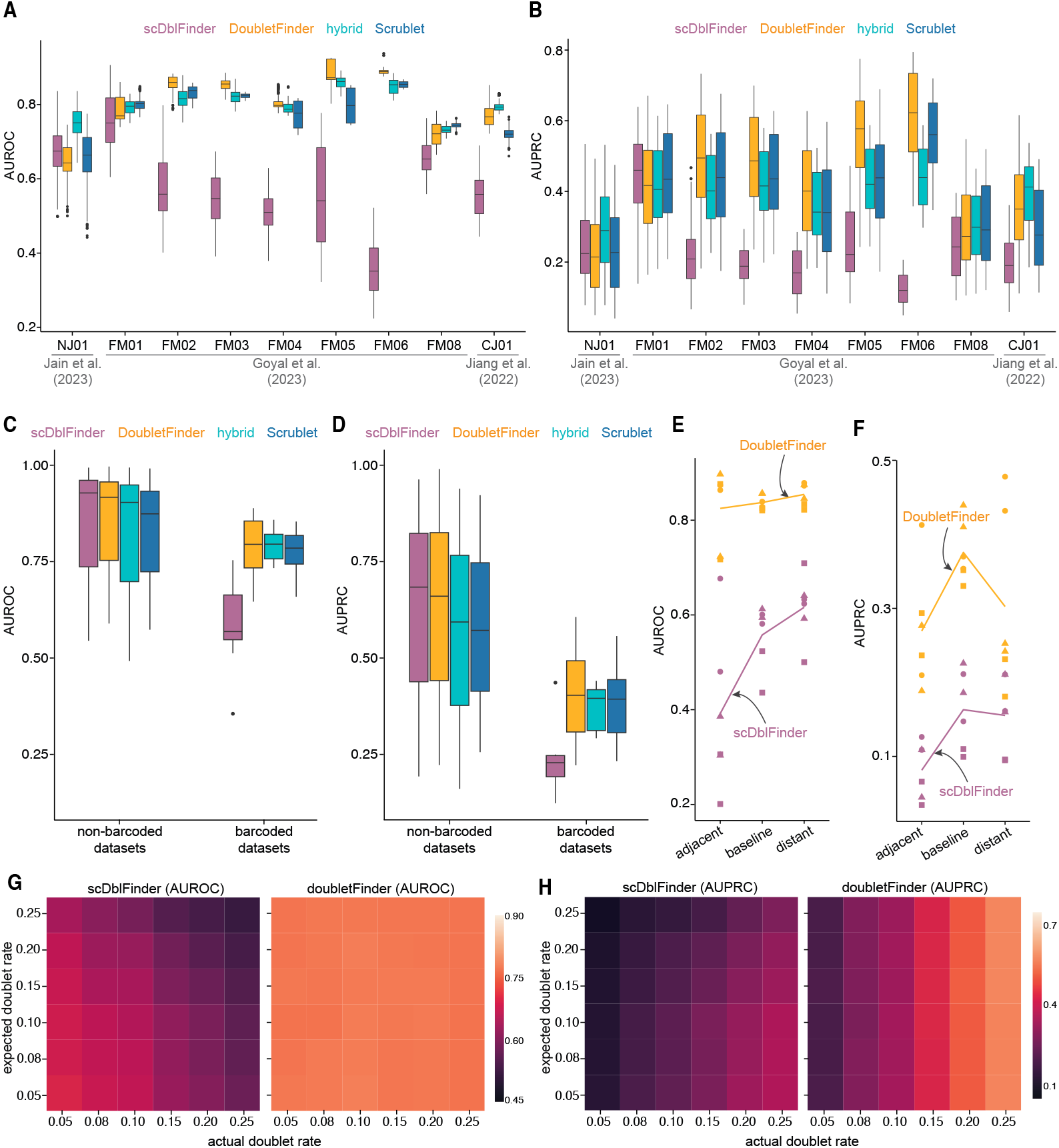
A. Color-coded boxplots of AUROC value for all four doublet detection methods. Each boxplot is calculated from the AUPOC value after running the respective method on each sample of the dataset three times with expected and actual doublet rate set to 0.05, 0.08, 0.1, 0.15, 0.2, and 0.25. B. Color-coded boxplots of AUPRC value for all four doublet detection methods calculated as panel A. C. Color-coded boxplots of AUROC value for four doublet detection methods on barcoded and non barcoded datasets. The boxplot for non-barcoded datasets was produced by running each dataset three times. The boxplot for barcoded datasets uses data from all barcoded datasets shown in panel A. D. Color-coded boxplots of AUPRC value for four doublet detection methods on barcoded and non barcoded datasets. The boxplot for non-barcoded datasets was produced by running each dataset three times. The boxplot for barcoded datasets uses data from all barcoded datasets shown in panel B. E. Scatter plot of AUROC score for three sets of datasets using DoubletFinder (yellow) and scDblFinder (pink). The line connects the mean of the datasets across the three conditions: adjacent, baseline, and distant. Each shape represents subsampled data from the particular experimental design (circle: FM02, triangle: FM03, and square: FM04). Cells were sampled from each experimental design twice. F. Scatter plot of AUPRC score for three sets of datasets using DoubletFinder (yellow) and scDblFinder (pink). The line connects the mean of the datasets across the three conditions: adjacent, baseline, and distant. Each shape represents subsampled data from the particular experimental design (circle: FM02, triangle: FM03, and square: FM04).Cells were sampled from each experimental design twice. G. Heatmap of AUROC average for all 27 tested FateMap barcoded datasets using scDblFinder and doubletFinder. scDblFinder AUROC ranges from 0.48 to 0.69. DoubletFinder AUROC ranges from 0.81 to 0.83. Tools are expected to have consistent AUROC regardless of expected or actual doublet rate. Heatmap ROC H. Heatmap of AUPRC average for all 27 tested FateMap barcoded datasets using scDblFinder and doubletFinder. scDblFinder AUPRC ranges from 0.13 to 0.36. DoubletFinder AUPRC ranges from 0.21 to 0.55. Tools are expected to have approximately consistent AUPRC value with a set actual doublet rate. AUPRC values are expected to increase as the actual doublet rate increase.Heatmap AUPRC

To evaluate doublet detection on these datasets, we selected four methods with varying performance from a recent benchmarking study [8]. DoubletFinder and scDblFinder, formerly known as doubletCells, were selected because they exhibited the highest and lowest accuracy, respectively [8]. Hybrid and Scrublet were selected because they showed strong performance despite using distinct underlying algorithms for predicting doublets [2, 5]. The benchmarking study primarily used area under Precision-Recall curve (AUPRC) and area under the receiver operating characteristic (AUROC) as their performance criteria. Note that AUROC is not necessarily representative of a doublet detection model’s performance in real datasets as true doublets often only account for a small percentage of the entire dataset [32]. AUPRC is a more robust metric for evaluating doublet detection methods given the inherent imbalance of singlet and doublet labels. An AUPRC value of 1 means the method can consistently identify doublets without mislabelling any singlets as doublets.

We evaluated the AUPRC and AUROC of the four methods on each of the barcoded datasets. Although we tested AUPRC and AUROC for a range of doublet rates (**Figure 2G-H** and **Supplementary 3D-E**), we present results for a reasonably practical true doublet rate of 8% (10X Genomics protocol predicts ∼8% doublet rate for maximum cell loading). The AUROC for all methods except scDblFinder in all barcoded datasets were fairly high, ranging from 0.6 to 0.85 (**Figure 2A**). In contrast, the AUPRC score for all four methods was consistently low (**Figure 2B**). The AUPRC values ranged from 0.5 to 0.8, with the overall average AUPRC being around 0.4 for all methods except scDblFinder, whose average AUPRC was around 0.23 (**Figure 2B**).

To directly compare the performance of doublet detection methods on barcoded and non-barcoded datasets, we selected three non-barcoded datasets [2,33,34] used in previous doublet detection algorithms for benchmarking comparisons. These datasets have annotated doublets and varying degrees of AUPRC and AUROC scores [8]. The AUROC values obtained from barcoded and non-barcoded datasets were consistent using all methods except scDblFinder (**Figure 2C** and [2]). However, the AUPRC values for the barcoded datasets were significantly lower than for the non-barcoded datasets, with scDblFinder having the greatest discrepancy (**Figure 2B,D**).

What is the basis of the difference in AUROC and AUPRC scores between our barcoded datasets and the non-barcoded datasets used in previous benchmarking studies? The datasets used for benchmarking tend to contain highly heterotypic doublets, meaning the scRNA-seq dataset is heterogeneous, often consisting of cells from different species or patients [8]. In contrast, the barcoded scRNA-seq datasets contain cells from a single system (and therefore often not as heterogeneous), a scenario more representative of many experimental designs. We wondered whether the low AUROC (for scDblFinder) and AUPRC scores are a result of relatively less heterogeneity. To answer this question, we sampled cells from adjacent (more similar), distal (less similar), or all (baseline) regions of the high-dimensional transcriptional space for a subset of datasets. Then, we calculated the AUPRC and AUROC scores for the strongest (DoubletFinder) and weakest (scDblFinder) performing methods (**Supplementary Figure 3A**, see **Methods**). We found a consistent increase in AUROC scores for both methods (especially scDblFinder) as cells become more transcriptionally heterogeneous (**Figure 2E**). The AUPRC scores were not as sensitive to the increasing variety of cells. The cells sampled from all regions did, however, have a slightly higher AUPRC score compared to adjacent or distal regions (**Figure 2F**). Collectively, the degree of transcriptional similarity between cells affects the AUROC scores, but not the AUPRC scores.

We questioned whether the AUPRC and AUROC scores for different methods were sensitive to the doublet rate, which can vary depending on the cell type and input cell numbers. We found that all methods except scDblFinder achieve consistent AUROC values regardless of expected or actual doublet rate (**Figure 2G** and **Supplementary Figure 3D**). In addition, we found that all tools achieve consistently higher AUPRC scores with increasing percentages of ground truth doublets, although the increase was modest for scDblFinder (**Figure 2H** and **Supplementary Figure 3E**). Furthermore, scDblFinder AUROC and AUPRC scores depended on the number of expected doublets, further undermining the robustness of scDblFinder in finding true singlets.

## Discussion

Here we develop and implement singletCode, a framework that leverages synthetic DNA barcoding prior to single-cell library prep reaction separation to extract “ground truth” singlets in scRNA-seq datasets. We test the feasibility of singleCode on a range of scRNA-seq datasets and evaluate the performance of doublet detection algorithms. We found comparatively low AUPRC and AUROC scores from different methods, particularly scDblFinder, for realistic scRNA-seq datasets not consisting of different species or patient samples. The AUPRC and AUROC scores were differentially sensitive to altering doublet fraction and degree of heterogeneity in scRNA-seq datasets. Our work underscores the importance of incorporating synthetic barcoding in experimental designs, especially when rare cell types are of particular interest. One limitation of our study is that we focused on one type of barcode technology (static and 100 base pairs). Future studies can extend singletCode to other types of synthetically introduced DNA barcoding technologies [23,25] and native cellular identifiers (e.g., mitochondrial mutations [35] or TCR sequences [10]).

Given the rapid rise in reported lineage barcoding datasets, singletCode provides a framework to identify ground truth singlets for downstream analysis. Alternatively, singletCode itself can be leveraged to test the performance of different doublet detection methods and choose an appropriate method given the data type. Particularly, we expect singletCode to be helpful for datasets consisting of rare or continuums of cell types (e.g., partial reprogramming, cell state transitions) and those which contain transcriptionally similar cells (e.g., clonally derived populations). Another application of singletCode could be to detect spatially adjacent or adhesive/sticky cells if the doublet barcoded cells tend to cluster within certain regions of the high dimensional transcriptional space. In principle, rich lineage barcoded datasets from different cell types and biological processes can be used to train deep learning models for broader querying of doublets on similar non-barcoded annotated atlases [36], including potential applications to spatial transcriptomics datasets. Together, singletCode illustrates the shortcomings of doublet detection methods and provides a generalizable and formalizable framework to identify true singlets.

## Methods

### Gini coefficient analysis for barcode uniformity and bias quantification

To perform the Gini analysis for the single cell data sets, we first simulated what the distribution of Gini coefficients would be from a uniform, exponential, and power distribution. We selected 20 random numbers from a given distribution and then calculated the Gini coefficient for those numbers. This process was then repeated approximately 20 times each for each distribution we simulated. For the power distribution, we used an alpha value of 1.5. Next, we preprocessed our single cell data by removing cells with feature counts fewer than the 10th percentile for that sample and cells with mitochondrial counts greater than the 60th percentile for that sample. Variable features were identified using a variance stabilizing transform for the top 2000 most variable genes. Neighbors were found using the top 50 dimensions, clusters were found using three resolutions (0.4, 0.8, 1.2), and UMAP projection was calculated using the top 50 dimensions.

For each of the three resolutions for a given sample, we calculated the Gini coefficient of the proportion of cells identified as singlets per cluster. We did this across multiple resolutions as they have differing amounts of clusters and we wanted to make sure the clustering of the data had no effect on the distribution of singlets.

### Nearest neighbor analysis for barcode uniformity and bias quantification

For the nearest neighbor analysis of the single-cell datasets, we first performed quality control steps, including thresholds for RNA counts, mitochondrial reads, and number of cells. Since the datasets cover several different cell types, the threshold values varied depending on the dataset, and we have provided them in **Supplementary Table 2**. To perform the nearest neighbor analysis, we extracted the top five nearest cell neighbors and the associated distances for each cell in the principal component space, and calculated the average neighbor distance. We performed this analysis on datasets composed either exclusively of singlets identified from the DNA lineage barcodes, or those that we randomly subsampled from the entire single-cell dataset consisting of singlets and multiplets. We performed random sampling and nearest neighbor calculation three times, ensuring the total subsampled cells to be the same number as the singlets for appropriate comparisons. The singlets’ average neighbor distance were normalized by subtracting the average neighbor distance of the subsampled cells. A mean value closer to zero indicates no preference for barcoding specific cells in high dimensional principal component spaces. A nonzero mean value suggests a preference for specific cells in high dimensional spaces or manifolds to be barcoded. Datasets displaying high standard deviations were affected by reduced cell counts attributed to high percentages of mitochondrial reads and a likely increase in cell death.

To account for what a “positive control” would look like for our nearest neighbor analysis, we artificially biased a control dataset by subsampling regions of principal component space from one of the samples (FM01_sample3, **Figure 1D**). We subsampled three restricted regions of the PCA plot by thresholding based on the PC1 vs PC2 plot (n = 1586 cells) to mimic highly clustered cells, and calculated the average nearest neighbor distance to compare them with the real experimental datasets.

### Generation of scRNA-seq datasets, singlets, and doublets

In order to simulate true doublets to most effectively benchmark the detection method performances, we identified true singlets by examining the mapping of the FateMap barcode to cell ID. Because FateMap barcodes are added to cells before the oil droplet encapsulates the cell, any cell ID which has only one associated FateMap barcode is a true singlet. To account for the scenario where multiple FateMap barcodes are added to each cell, we further characterized all cell IDs that are associated with the same combination of FateMap barcodes as singlets.

Lastly, it is possible that some cells receive ambient FateMap barcodes. This should result in one of such FateMap barcode having a significantly higher UMI count since all other barcodes are ambient FateMap barcodes for a given cell. We therefore also consider these cells as singlets. There are two cases where cells may be classified as such a singlet. First, the cell has only one FateMap barcode with at least 10 associated UMI. Alternatively, it could have multiple FateMap barcodes with at least 10 associated UMI, but one of these FateMap barcodes has at least 50 more UMI than the median of all FateMap UMI count for that cell. We used these three strategies to label true singlets in the dataset.

To simulate the doublets, we randomly selected the count data from two cells we identify as true singlets. We then averaged the counts from these two cells to generate simulated doublets as performed previously [1]. The final scRNA-seq datasets are generated by adding such simulated doublets into the datasets at different percentages (5-25%).

### Benchmark Environment and Parameter Settings

We ran each doublet detection method three times on each scRNA-seq dataset with expected and actual doublet rate set to 0.05, 0.08, 0.1, 0.15, 0.2, and 0.25. This means each method is run 125 times in total per dataset. The dataset is loaded with Seurat and converted to *SingleCellExperiment* in R [37] if necessary. All algorithms were run with recommended settings following their official tutorial.

DoubletCells (scDblFinder): This method was executed in R following the instructions at https://bioconductor.org/packages/release/bioc/vignettes/scDblFinder/inst/doc/scDblFinder.html

DoubletFinder: This method was executed in R following the instructions at https://github.com/chris-mcginnis-ucsf/DoubletFinder

Scrublet: This method was executed in Python following the instructions at https://github.com/AllonKleinLab/scrublet/blob/master/examples/scrublet_basics.ipynb

Hybrid: This method was executed in R following the instructions at https://github.com/kostkalab/scds

To evaluate how DoubletFinder and scDblFinder performs on adjacent, baseline, and distal data, we further selected FM02, FM03, FM04 and performed standard Seurat processing pipeline on them to generate cluster assignment as well as the corresponding UMAP projection. We defined three sets of datasets: baseline, which consists of 2,000 cells randomly subsampled from each dataset; adjacent, which consists of cells from neighboring clusters; and distal, which consists of cells from distal clusters. We then used the same settings to annotate doublets and calculated AUROC and AUPRC with both tools.

## Acknowledgements

We thank members of the Goyal lab for helpful discussions, especially Ian Mellis who discussed this project with YG during its inception. YG acknowledges support from Northwestern University’s startup funds and the Burroughs Wellcome Fund Career Awards at the Scientific Interface. ZZ and MEM were supported by funds from YG. KK acknowledges support from the University of Pennsylvania Medical Scientist Training Program.

## Code and data availability

All code and data developed in this manuscript is provided at: https://github.com/GoyalLab/fatemap_multiplet_public. All raw data was taken from published manuscripts (see Gene Expression Omnibus accession GSE233766, GSE198729, GSE227151 for scRNA-seq datasets; find barcoding datasets at Figshare (https://doi.org/10.6084/m9.figshare.22798952; https://doi.org/10.6084/m9.figshare.22802888; https://doi.org/10.6084/m9.figshare.19126985; https://doi.org/10.6084/m9.figshare.22251223.v1, https://doi.org/10.6084/m9.figshare.22236949.v1, https://doi.org/10.6084/m9.figshare.22236955.v1).

## Author Contributions

YG conceived and designed the project. ZZ designed, performed, and analyzed the doublet algorithm performance supervised by YG. KK and MEM performed singlet uniformity analysis. MEM and YG organized the main and supplementary figures and tables. YG, ZZ, and MEM wrote the manuscript with input from KK.

## Figures

**Supplementary Figure 1**.

A. Example UMAP projection representation of one of the samples: FM01 sample 2. Singlets are evenly distributed across UMAP space and are not overrepresented in any one cluster.

B. The average of the 5 nearest neighbor distances for singlets in each sample, including an artificially biased control based on subsampling FM01 sample 3. Samples for FM08 and NJ01 are grouped due to low cell counts. Error bars represent standard deviation and the dot represents a sample’s mean nearest neighbor distance normalized to the average neighbor distance for the subsampled total cells (horizontal line represents a zero value).

**Supplementary Figure 2**.

A.(top) UMAP projection showing clustering across three cluster resolutions (0.4, 0.8, 1.2). (bottom) Corresponding proportion of singlets per cluster across those clustering resolutions.

B. Violin plots of distribution of QC metrics (number of features, number of counts, percent mitochondrial counts) for cluster resolution of 0.4.

C. Violin plots of distribution of QC metrics (number of features, number of counts, percent mitochondrial counts) for cluster resolution of 0.8.

D. Violin plots of distribution of QC metrics (number of features, number of counts, percent mitochondrial counts) for cluster resolution of 1.2.

**Supplementary Figure 3**.

A. UMAP projections highlighted by cells selected to form distal, adjacent, and baseline datasets used for assessing the impact of cell similarity on doublet detection performance for FM02.

B. UMAP projections highlighted by cells selected to form distal, adjacent, and baseline datasets used for assessing the impact of cell similarity on doublet detection performance for FM03.

C. UMAP projections highlighted by cells selected to form distal, adjacent, and baseline datasets used for assessing the impact of cell similarity on doublet detection performance for FM04.

D. Heatmap of AUROC average for all 27 tested FateMap barcoded datasets using hybrid and scrublet. Hybrid AUROC ranges from 0.79 to 0.83. Scrublet AUROC is consistently around 0.80. Tools are expected to have consistent AUROC regardless of expected or actual doublet rate.

E. Heatmap of AUPRC average for all 27 tested FateMap barcoded datasets using hybrid and scrublet. Hybrid AUPRC ranges from 0.24 to 0.54. Scrublet AUPRC ranges from 0.24 to 0.58. Tools are expected to have approximately consistent AUPRC value with a set actual doublet rate. AUPRC values are expected to increase as the actual doublet rate increases.

**Supplementary Table 1**. (link)

Specifications for barcoded and non-barcoded datasets used in this analysis including origin, treatment, and cell/singlet counts.

**Supplementary Table 2**. (link)

Neighbor analysis quality control thresholds for all barcoded datasets. Quality control parameters include minimum and maximum RNA count, and maximum percentage of mitochondrial counts.

## References

1. Bernstein NJ, Fong NL, Lam I, Roy MA, Hendrickson DG, Kelley DR. Solo: Doublet Identification in Single-Cell RNA-Seq via Semi-Supervised Deep Learning. Cell Syst. 2020;11:95–101.e5.

2. Xi NM, Li JJ. Benchmarking Computational Doublet-Detection Methods for Single-Cell RNA Sequencing Data. Cell Syst. 2021;12:176–94.e6.

3. Luecken MD, Theis FJ. Current best practices in single-cell RNA-seq analysis: a tutorial. Mol Syst Biol. 2019;15:e8746.

4. McGinnis CS, Murrow LM, Gartner ZJ. DoubletFinder: Doublet Detection in Single-Cell RNA Sequencing Data Using Artificial Nearest Neighbors. Cell Syst. 2019;8:329–37.e4.

5. Wolock SL, Lopez R, Klein AM. Scrublet: Computational Identification of Cell Doublets in Single-Cell Transcriptomic Data. Cell Syst. 2019;8:281–91.e9.

6. DePasquale EAK, Schnell DJ, Van Camp P-J, Valiente-Alandí Í, Blaxall BC, Grimes HL, et al. DoubletDecon: Deconvoluting Doublets from Single-Cell RNA-Sequencing Data. Cell Rep. 2019;29:1718–27.e8.

7. Lun ATL, McCarthy DJ, Marioni JC. A step-by-step workflow for low-level analysis of single-cell RNA-seq data with Bioconductor. F1000Res. 2016;5:2122.

8. Bais AS, Kostka D. scds: computational annotation of doublets in single-cell RNA sequencing data. Bioinformatics. 2020;36:1150–8.

9. McGinnis CS, Patterson DM, Winkler J, Conrad DN, Hein MY, Srivastava V, et al. MULTI-seq: sample multiplexing for single-cell RNA sequencing using lipid-tagged indices. Nat Methods. 2019;16:619–26.

10. Sun B, Bugarin-Estrada E, Overend LE, Walker CE, Tucci FA, Bashford-Rogers RJM. Double-jeopardy: scRNA-seq doublet/multiplet detection using multi-omic profiling. Cell Rep Methods. 2021;1:None.

11. Stoeckius M, Zheng S, Houck-Loomis B, Hao S, Yeung BZ, Mauck WM 3rd, et al. Cell Hashing with barcoded antibodies enables multiplexing and doublet detection for single cell genomics. Genome Biol. 2018;19:224.

12. Bhang H-EC, Ruddy DA, Krishnamurthy Radhakrishna V, Caushi JX, Zhao R, Hims MM, et al. Studying clonal dynamics in response to cancer therapy using high-complexity barcoding. Nat Med. 2015;21:440–8.

13. Biddy BA, Kong W, Kamimoto K, Guo C, Waye SE, Sun T, et al. Single-cell mapping of lineage and identity in direct reprogramming. Nature. 2018;564:219–24.

14. Weinreb C, Rodriguez-Fraticelli A, Camargo F, Klein AM. Lineage tracing on transcriptional landscapes links state to fate during differentiation [Internet]. bioRxiv. 2018 [cited 2019 Oct 4]. p. 467886. Available from: https://www.biorxiv.org/content/10.1101/467886v2

15. Gutierrez C, Al’Khafaji AM, Brenner E, Johnson KE, Gohil SH, Lin Z, et al. Multifunctional barcoding with ClonMapper enables high-resolution study of clonal dynamics during tumor evolution and treatment. Nature Cancer. 2021;2:758–72.

16. Oren Y, Tsabar M, Cuoco MS, Amir-Zilberstein L, Cabanos HF, Hütter J-C, et al. Cycling cancer persister cells arise from lineages with distinct programs. Nature. 2021;596:576–82.

17. Frieda KL, Linton JM, Hormoz S, Choi J, Chow K-HK, Singer ZS, et al. Synthetic recording and in situ readout of lineage information in single cells. Nature. 2017;541:107–11.

18. Umkehrer C, Holstein F, Formenti L, Jude J, Froussios K, Neumann T, et al. Isolating live cell clones from barcoded populations using CRISPRa-inducible reporters. Nat Biotechnol. 2021;39:174–8.

19. Emert BL, Cote CJ, Torre EA, Dardani IP, Jiang CL, Jain N, et al. Variability within rare cell states enables multiple paths toward drug resistance. Nat Biotechnol. 2021;39:865–76.

20. Tian L, Tomei S, Schreuder J, Weber TS, Amann-Zalcenstein D, Lin DS, et al. Clonal multi-omics reveals Bcor as a negative regulator of emergency dendritic cell development. Immunity. 2021;54:1338–51.e9.

21. Leighton J, Hu M, Sei E, Meric-Bernstam F, Navin NE. Reconstructing mutational lineages in breast cancer by multi-patient-targeted single cell DNA sequencing [Internet]. bioRxiv. 2021 [cited 2021 Dec 4]. p. 2021.11.16.468877. Available from: https://www.biorxiv.org/content/10.1101/2021.11.16.468877v1

22. Rodriguez-Fraticelli AE, Weinreb C, Wang S-W, Migueles RP, Jankovic M, Usart M, et al. Single-cell lineage tracing unveils a role for TCF15 in haematopoiesis. Nature. 2020;583:585–9.

23. Pillai M, Hojel E, Jolly MK, Goyal Y. Unraveling non-genetic heterogeneity in cancer with dynamical models and computational tools. Nature Computational Science [Internet]. 2023; Available from: https://www.nature.com/articles/s43588-023-00427-0

24. Fennell KA, Vassiliadis D, Lam EYN, Martelotto LG, Balic JJ, Hollizeck S, et al. Non-genetic determinants of malignant clonal fitness at single-cell resolution. Nature. 2022;601:125–31.

25. Sankaran VG, Weissman JS, Zon LI. Cellular barcoding to decipher clonal dynamics in disease. Science. 2022;378:eabm5874.

26. Goyal Y, Busch GT, Pillai M, Li J, Boe RH, Grody EI, et al. Diverse clonal fates emerge upon drug treatment of homogeneous cancer cells. Nature [Internet]. 2023; Available from: http://dx.doi.org/10.1038/s41586-023-06342-8

27. Jain N, Goyal Y, Dunagin MC, Cote CJ, Mellis IA, Emert B, et al. Retrospective identification of intrinsic factors that mark pluripotency potential in rare somatic cells [Internet]. bioRxiv. 2023 [cited 2023 Mar 7]. p. 2023.02.10.527870. Available from: https://www.biorxiv.org/content/10.1101/2023.02.10.527870v1

28. Jiang CL, Goyal Y, Jain N, Wang Q, Truitt RE, Coté AJ, et al. Cell type determination for cardiac differentiation occurs soon after seeding of human-induced pluripotent stem cells. Genome Biol. 2022;23:90.

29. Reffsin S, Miller J, Ayyanathan K, Dunagin MC, Jain N, Schultz DC, et al. Single cell susceptibility to SARS-CoV-2 infection is driven by variable cell states [Internet]. bioRxiv. 2023 [cited 2023 Jul 22]. p. 2023.07.06.547955. Available from: https://www.biorxiv.org/content/10.1101/2023.07.06.547955v1.abstract

30. Weinreb C, Rodriguez-Fraticelli A, Camargo FD, Klein AM. Lineage tracing on transcriptional landscapes links state to fate during differentiation. Science [Internet]. 2020;367. Available from: http://dx.doi.org/10.1126/science.aaw3381

31. Schuh L, Saint-Antoine M, Sanford EM, Emert BL, Singh A, Marr C, et al. Gene Networks with Transcriptional Bursting Recapitulate Rare Transient Coordinated High Expression States in Cancer. Cell Syst. 2020;10:363–78.e12.

32. Alexandari AM, Kundaje A, Shrikumar A. A General Framework for Abstention Under Label Shift [Internet]. arXiv [stat.ML]. 2018. Available from: http://arxiv.org/abs/1802.07024

33. Zheng GXY, Terry JM, Belgrader P, Ryvkin P, Bent ZW, Wilson R, et al. Massively parallel digital transcriptional profiling of single cells. Nat Commun. 2017;8:14049.

34. Kang HM, Subramaniam M, Targ S, Nguyen M, Maliskova L, McCarthy E, et al. Multiplexed droplet single-cell RNA-sequencing using natural genetic variation. Nat Biotechnol. 2018;36:89–94.

35. Miller TE, Lareau CA, Verga JA, DePasquale EAK, Liu V, Ssozi D, et al. Mitochondrial variant enrichment from high-throughput single-cell RNA sequencing resolves clonal populations. Nat Biotechnol [Internet]. 2022; Available from: http://dx.doi.org/10.1038/s41587-022-01210-8

36. Heimberg G, Kuo T, DePianto D, Heigl T, Diamant N, Salem O, et al. Scalable querying of human cell atlases via a foundational model reveals commonalities across fibrosis-associated macrophages [Internet]. bioRxiv. 2023 [cited 2023 Aug 5]. p. 2023.07.18.549537. Available from: https://www.biorxiv.org/content/10.1101/2023.07.18.549537v1

37. Amezquita RA, Lun ATL, Becht E, Carey VJ, Carpp LN, Geistlinger L, et al. Orchestrating single-cell analysis with Bioconductor. Nat Methods. 2020;17:137–45.

