## Supplementary figures and images for "singletCode: synthetic barcodes identify singlets in scRNA-seq datasets and evaluate doublet algorithms"

### Supplementary Figure 1

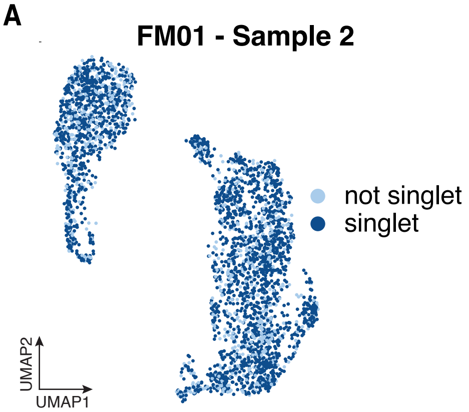

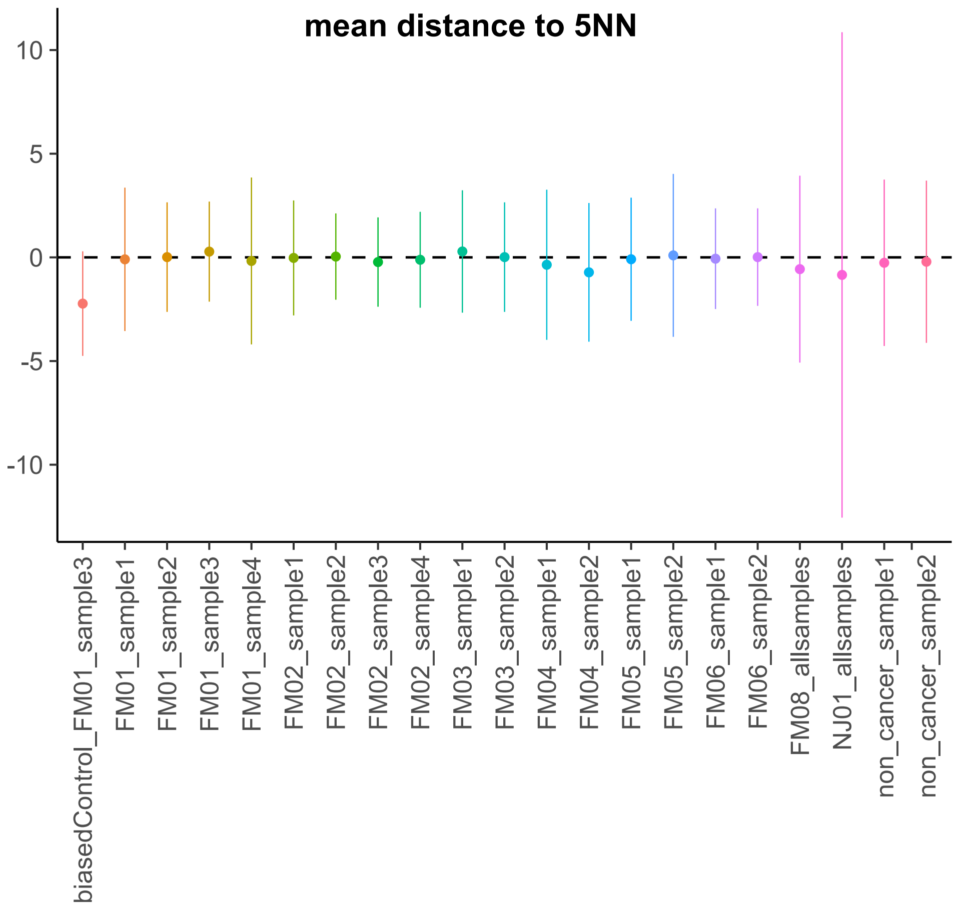

### Supplementary Figure 2

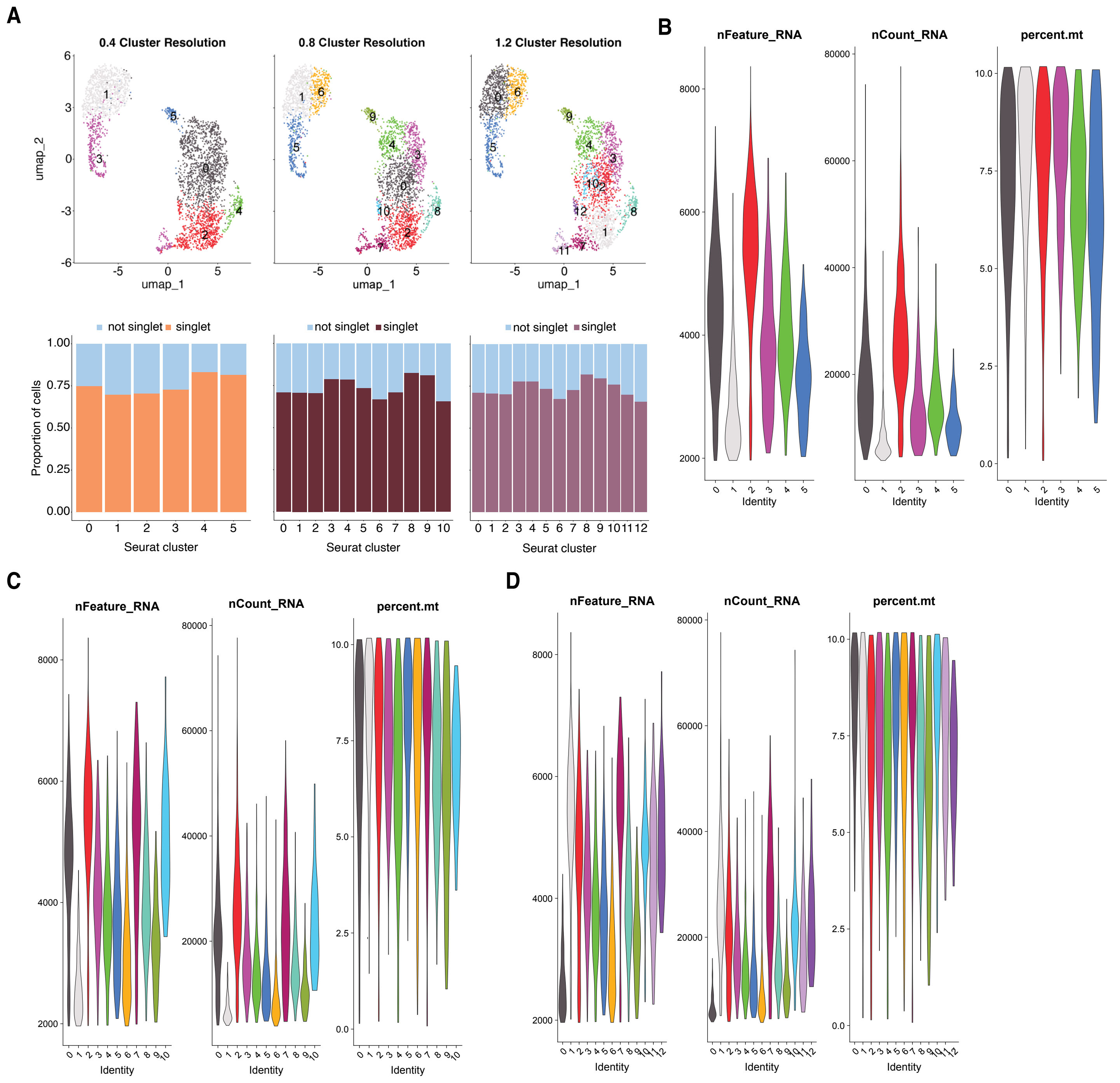

### Supplementary Figure 3

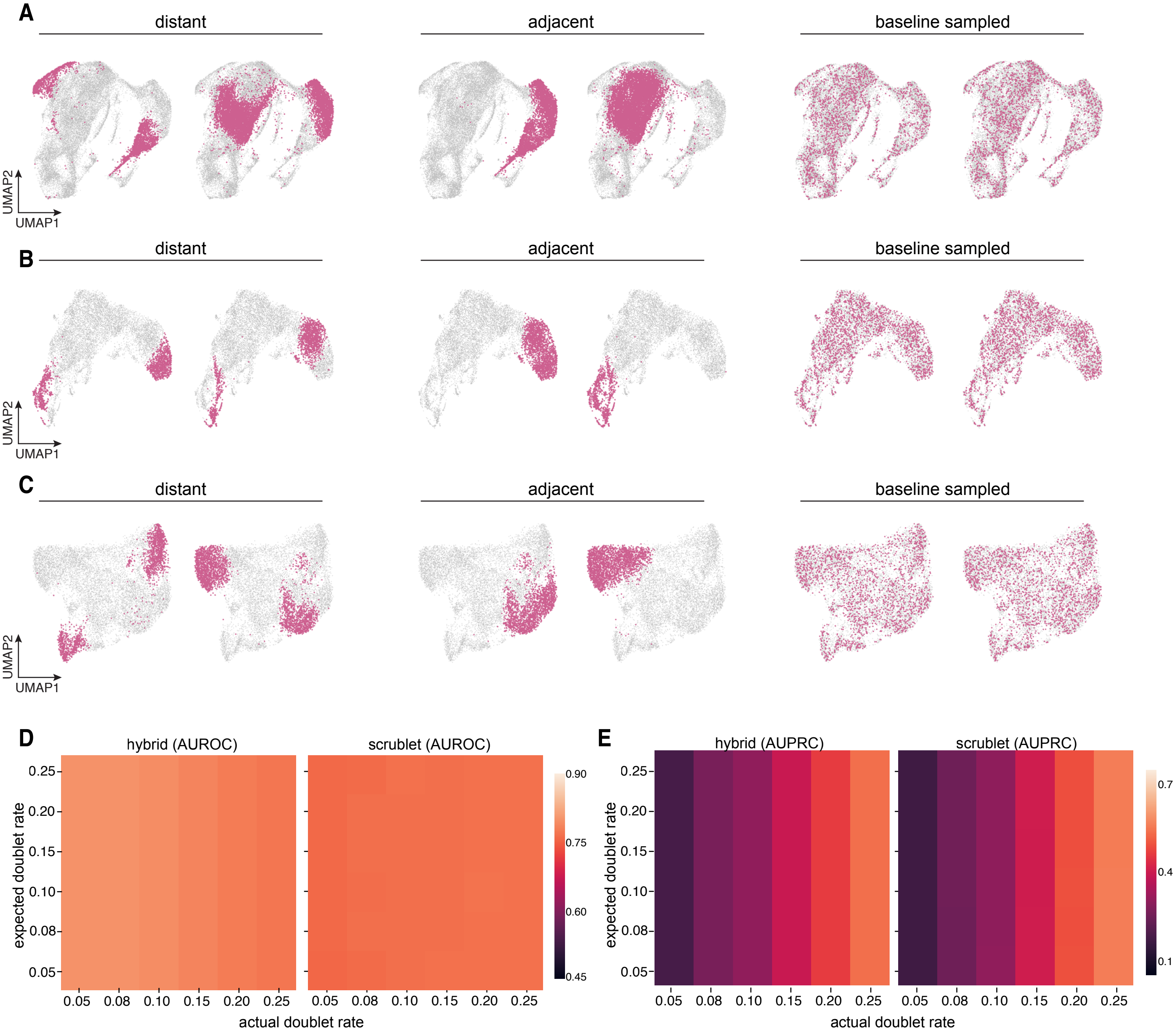
