## Supplementary Table 1 for "singletCode: synthetic barcodes identify singlets in scRNA-seq datasets and evaluate doublet algorithms"

**Supplementary Table 1. Specifications for barcoded and non-barcoded datasets**

| Alias ID | Sample type | Organism | Tissue | Cell line | Genotype | Treatment | Total samples | Total cells | Total singlets | Publication |
| --- | --- | --- | --- | --- | --- | --- | --- | --- | --- | --- |
| FM01 | Melanoma patient resistant cell line | Homo sapiens | Skin-derived | WM989 A6-G3 | BRAF – V600E | Targeted therapy | 4 | 25570 | 20477 | Goyal et al., *Nature*, 2023 |
| FM02 | Melanoma patient resistant cell line | Homo sapiens | Skin-derived | WM989 A6-G3 | BRAF – V600E | Targeted therapy | 4 | 28450 | 23841 | Goyal et al., *Nature*, 2023 |
| FM03 | Melanoma patient resistant cell line | Homo sapiens | Skin-derived | WM989 A6-G3 | BRAF – V600E | Targeted therapy | 2 | 10107 | 8841 | Goyal et al., *Nature*, 2023 |
| FM04 | Breast cancer patient resistant cell line | Homo sapiens | Breast; mammary gland | MDA-MB-231-D4 | WT | Chemotherapy, cytotoxic | 2 | 13220 | 10136 | Goyal et al., *Nature*, 2023 |
| FM05 | Melanoma patient resistant cell line | Homo sapiens | Skin-derived | WM989 A6-G3 | BRAF – V600E | Targeted therapy | 2 | 33655 | 26268 | Goyal et al., *Nature*, 2023 |
| FM06 | Melanoma patient untreated cell line | Homo sapiens | Skin-derived | WM989 A6-G3 | BRAF – V600E | No treatment | 2 | 15509 | 13032 | Goyal et al., *Nature*, 2023 |
| FM08 | Primary melanocytes | Homo sapiens | Skin-derived | FOM230-1 | WT | No treatment | 3 | 5074 | 4851 | Goyal et al., *Nature*, 2023 |
| non-cancer | HIPS differentiation | Homo sapiens | PBMCs | PENN123i-SV20 | WT | Cardiac differentiation signal | 2 | 5558 | 3453 | Jiang et al., *Genome Biol*, 2022 |
| NJ01 | Stem cell reprogramming | Homo sapiens | Fibroblasts | hiF-T | WT | OKSM | 6 | 3901 | 3610 | Jain et al., *bioRxiv,* 2023 |
| non- barcoded: | | | | | | | | | | |
| hm-12k | Synthetic dataset | Homo sapiens, Mus musculus | Human kidney, Murine fibroblasts | HEK293T, NIH3T3 | WT | No treatment | 2 | 12820 | 12090 | Zhang et al., *Nat Commun*, 2017 |
| J293t | Potentially faulty annotation | Homo sapiens | Human kidney, T-lymphocytes | HEK293T, Jurkat | WT | No treatment | 1 | 500 | 458 | Kang et al., *Nat Biotechnol,* 2018 |
| pbmc-2ctrl-dm | Patient PBMCs | Homo sapiens | Systemic lupus erythematosus (SLE) PBMCs | Patient- derived | Patient- derived | No treatment | 2 | 13913 | 12315 | Stoeckius et al., *Genome Biol*, 2018 |
