## Supplementary Table 2 for "singletCode: synthetic barcodes identify singlets in scRNA-seq datasets and evaluate doublet algorithms"

**Supplementary Table 2: Neighbor analysis quality control thresholds**

| Dataset | **Minimum nCount.RNA** | **Maximum nCount.RNA** | **Maximum percent.mt** |
| --- | --- | --- | --- |
| FM01 | 200 | 6000 | 20 |
| FM02 | 200 | 6000 | 20 |
| FM03 | 200 | 6000 | 40 |
| FM03 biased | 200 | 6000 | 20 |
| FM04 | 200 | 6000 | 40 |
| FM05 | 200 | 6000 | 50 |
| FM06 | 200 | 6000 | 20 |
| FM08 | 200 | 10000 | 40 |
| NJ01 | 200 | 8000 | 21 |
| non-cancer | 200 | 7500 | 25 |
